# Analysis of genomic DNA methylation and gene transcription modulation induced by *DNMT3A* deficiency in HEK293 cells

**DOI:** 10.1101/2020.07.11.198895

**Authors:** Mengxiao Zhang, Jiaxian Wang, Qiuxiang Tian, Lei Feng, Hui Yang, Nan Zhu, Xingchen Pan, Jianwei Zhu, Peng Chen, Huili Lu

## Abstract

DNA methylation is an important epigenetic modification associated with transcriptional repression, and plays key roles in normal cell growth as well as oncogenesis. Among the three main DNA methyltransferases (DNMT1, DNMT3A, and DNMT3B), DNMT3A mediates *de novo* DNA methylation with partial functional redundancy with DNMT3B. However, the general effects of DNMT3A and its downstream gene regulation profile are yet to be unveiled. In the present study, we used CRISPR/Cas9 technology to successfully create DNMT3A deficient human embryonic kidney cell line HEK293, with frameshift mutations in its catalytic domain. Our results showed that the cell growth slowed down in *DNMT3A* knockout cells. UPLC-MS analysis of DNMT3A deficient cells showed that the genome-level DNA methylation was reduced by 21.5% and led to an impairment of cell proliferation as well as a blockage of MAPK and PI3K-Akt pathways. Whole genome RNA-seq revealed that *DNMT3A* knockout up-regulated expression of genes and pathways related to cell metabolism but down-regulated those involved in ribosome function, which explained the inhibition of cell growth and related signal pathways. Further, bisulfite DNA treatment showed that *DNMT3A* ablation reduced the methylation level of *DNMT3B* gene as well, indicating the higher DNMT3B activity and thereby explaining those down-regulated profiles of genes.

## 1 INTRODUCTION

DNA methylation is an epigenetic modification with widespread effects on gene expression. High levels of promoter DNA methylation is usually associated with gene silencing (You & Jones, 2012). Abnormal DNA methylation is involved in the development of multiple malignancies, such as solid tumors and leukemia (Banaszak et al., 2018; Fan et al., 2010; Gao et al., 2013; Lee et al., 2005; Mirza et al., 2013; Montgomery et al., 2004; Yang et al., 2016). In vertebrates, cytosine methylation on CpG dinucleotides is the predominant form of methylation catalyzed by DNA methyltransferase 1 (DNMT1) (Bestor, 1992) and established *de novo* by DNMT3A and DNMT3B (Okano et al., 1999; Okano et al., 1998).

To investigate the underlying mechanisms responsible for locus-specific or global methylation, *in vivo* and *in vitro* models of DNMTs deficiency have been developed (Huang et al., 2017). In mice, knockout of *DNMT1* or *DNMT3B* can cause early embryo death. In contrast, *DNMT3A* knockout mice can be born normally but develop developmental defects and die premature soon after birth (Okano et al., 1999). These observations highlight that DNMT3A plays specific roles in regulating chromatin methylation during the development after birth (Okano et al., 1999; Riggs & Xiong, 2004). Similarly, in human embryonic cells, individual or simultaneous disruption of DNMT3A or DNMT3B resulted in viable, pluripotent cell lines, but deletion of DNMT1 resulted in rapid cell death (Liao et al., 2015). Banaszak et al. mutated DNMT3A in K562 leukemia cells and the derived cell lines showed impaired cell growth (Banaszak et al., 2018). Although almost all cells can survive DNMT3A mutation, reports have shown paradoxical hyper-methylation of genes, or no changes in global or regional DNA methylation patterns in response to DNMT3A knockdown (Banaszak et al., 2018; Challen et al., 2012). Hence, the exact roles of DNMT3A are yet to be elucidated.

CRISPR/Cas9 system is an efficient genome editing technique developed in recent years (Horvath & Barrangou, 2010). Comparing with the traditional knockout techniques, such as zinc finger nuclease technology (ZFN) and transcriptional activation effect factor nuclease technology (TALEN), CRISPR/Cas9 is comparatively easy to implement, is cost and time-effective, as well as has higher efficiency. CRISPR/Cas9 technique has been successfully used in human cells, zebrafish, mice, and bacterial genome modification (Le Cong et al., 2013; Mali et al., 2013). In the present study, we used CRISPR/Cas9 technology to establish a *DNMT3A* knockout cell line derived from HEK293T, a human embryonic kidney cell line. We performed detailed transcriptomic and epigenetic analyses, in addition to physiology measurements, to discover the impact of DNMT3A deficiency on cell proliferation and metabolism, as well as to identify genes which are potentially regulated by DNMT3A.

## 2 MATERIALS AND METHODS

### 2.1 Cell culture and reagents

Wild *type* HEK293 cells (HEK293T) were obtained from the Type Culture Collection of the Chinese Academy of Sciences (Shanghai, China) and detected to be negative for mycoplasma contamination using the Myco-Blue mycoplasma detector (Vazyme; Nanjing, Jiangsu, China). Cells were cultured in high glucose DMEM supplemented with 10% FBS, incubated at 37°C with 5% CO_2_ in a humidified cell incubator (Thermo Fisher Scientific; OH, USA). The plasmid pX330 carrying CRISPR/Cas9 system was kindly provided by Dr. Feng Zhang (MIT) (L. Cong et al., 2013). Competent cells of the *E. coli* strains DH5α were purchased from Microgene (Shanghai, China). All media and supplements were purchased from Gibco (Thermo Fisher Scientific; Waltham, MA, USA). Cell growth and viability were monitored with a cell counter (Countstar; Shanghai, China).

### 2.2 SgRNA design and DNMT3A disruptive vector construction

Two sgRNAs targeting exon 19 of *DNMT3A* (GeneBank ID 806904736) were designed using the web tool provided by Dr. Zhang’s lab (http://crispr.mit.edu) as shown in **Fig. 1**. To construct the sgRNA plasmids, single strand primers were designed and synthesized as sgRNA1-forward: 5’-CACCGCATGATGCGCGGCCCAAGG-3’, sgRNA1-reverse 5’-AAACCCTTGGGCCGCGCATCATGC-3’, sgRNA2-forward 5’-CACCGCTCACTAATGGCTTCTACCT-3’ and sgRNA1-reverse 5’-AAACAGGTAGAAGCCATTAGTGAGC-3’. Each pair of primers were annealed to generate double-stranded cDNA, phosphorylated by T4 polynucleotide kinase at the 5’ ends (NEB, Ipswich, MA) at 37°C for 30 min, and further ligated into BbsI digested pX330 plasmids by T4 DNA ligase (Takara; Kusatsu, Shiga, Japan). The ligate was transformed to DH5α competent cells for culture overnight. Then the grown clones were selected for sequencing to get the right constructed plasmids pX330-sgRNA1 and pX330-sgRNA2.

**Figure 1.**
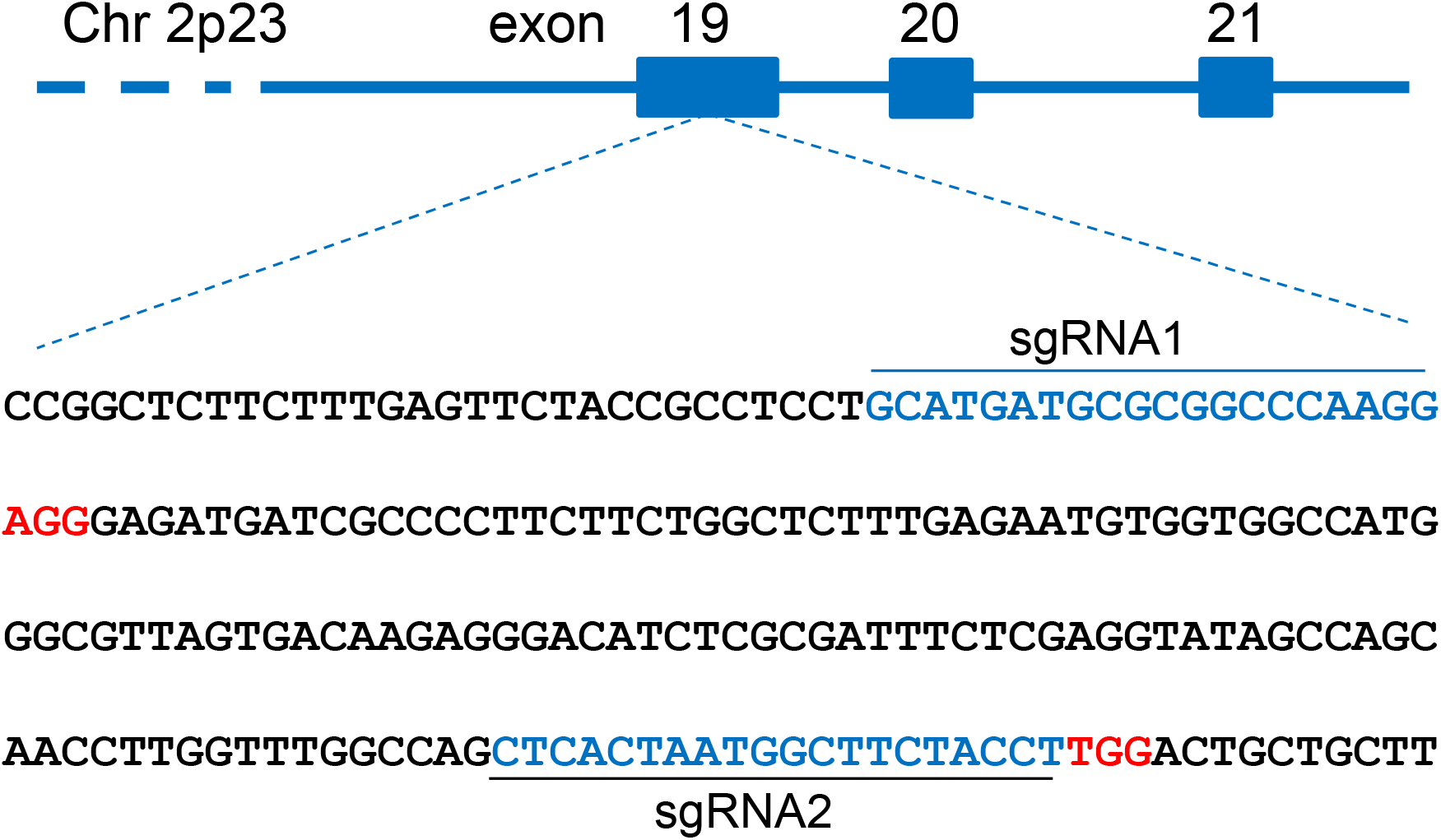
Location and design of sgRNAs. The sgRNA sequences are shown in blue, and protospacer adjacent motif (PAM) bases are in red.

### 2.3 Transfection of HEK293 cells

HEK293 cells were seeded at 2×10^5^ cells/well into 12-well plate one day prior to transfection. When reached 70-80% confluence, the cells were co-transfected with pX330-sgRNA1 and pX330-sgRNA2 at a molar ratio of 1:1, since it was reported that double sgRNAs could result in higher editing efficiency than single one (Zheng et al., 2014). The transfection was performed using Lipofectamine 2000 reagent (Invitrogen, CA, USA) according to manufacturer’s instructions.

### 2.4 DNMT3A knockout clones selection

HEK293 cell pool transfected with pX330-sgRNAs were seeded into 96-well plates at the density of 0.5 cell per well for limiting dilution. After about ten days’ incubation, the plates were examined for single cell clones under microscope. When grew to about 80% confluent in the well, the clones would be detached for subpopulation and the genomic DNA was extracted with *QuickExtract* DNA extraction solution (Epicenter; MD, USA) for PCR verification, using primers HEK293-DNMT3A-For (5’-GTACCATCCTGTCCCCTCCAC-3’) and HEK293-DNMT3A-Rev (5’-GGCTCAGGGTTAAACGGGGA-3’), which can amplify a 798 bp fragment for HEK293 wild-type cells. By sequencing the amplified fragments, the clone with disrupted DNMT3A was selected and designated to be DNMT3A KO cell line.

### 2.5 DNMT3A knockout cells proliferation curve

DNMT3A KO and WT cells were cultured and seeded at 3×10^4^ cells/well into 12-well plate. Cells were counted every 24 h for consecutive 6 days. And cell proliferation curves were compared between the two cell lines.

### 2.6 Western blot analysis

*DNMT3A* KO and WT HEK293 cells of 1 ×10^6^ were washed with PBS, lysed using 100 μL RAPA lysis buffer containing protease inhibitors cocktail (Roche; Penzberg, Germany), and separated by a 10% SDS-PAGE. After transferring onto a 0.45 μm PVDF membranes, immunoblotting was performed. For detection of DNMT3A deficiency, primary mouse monoclonal antibody against GAPDH (Sangon; Shanghai, China) and polyclonal rabbit-anti-human DNMT3A (Sangon) were used at 1:1000 dilution. For detection of MAPK and PI3K-Akt pathways, primary monoclonal antibodies against human Erk (137F5; Cell Signaling Technology; Danvers, MA, USA), phosphor-Erk (197G2; Cell Signaling Technology), JNK (D-2; Santa Cruz; Dallas, TX, USA), phosphor-JNK (G9; Cell Signaling Technology), Akt (11E7; Cell Signaling Technology), and phosphor-Akt (244F9; Cell Signaling Technology) were used. HRP-conjugated anti-mouse IgG or anti-rabbit IgG antibodies (Jackson ImmunoResearch; PA, USA) were used for secondary antibodies. Signals were detected with enhanced chemiluminescence (Millipore; MA, USA) and visualized with a gel imaging system (Tanon; Shanghai, China).

### 2.7 Genome-wide DNA methylation analysis by UPLC-ESI-MS/MS

Genomic DNA of cells were extracted by AxyPrep Kit (Axygen; Hangzhou, Zhejiang, China) and RNase A was added to remove RNA. Then the genomic DNA was hydrolyzed by DNase I at 37 °C for 1 h, denatured at 100 °C for 3 min, and immediately cooled down on ice for 10 min, then treated with *Nuclease* P1 at 37°C for 16 h, followed by treatment of alkaline phosphatase at 37 °C for 2 h. The nucleotides were stored at −20 °C before UPLC-ESI-MS-MS detection.

Acquity UPLC (Waters, USA) coupled with Triple Quad™ 5500 mass spectrometry (Sciex, USA) was used to quantitatively analyze m^5^dC and dG. UPLC-ESI-MS/MS method was established to evaluate DNA methylation status of genome (Putra et al., 2014). Reference nucleotide standards of A, G, T, C, dA, dG, dC, U and m^5^dC were purchased from Sigma (Sigma Aldrich, St. Louis, MO, USA) and dissolved in H_2_O to a final concentration of 1.0 mg/ml. UPLC and electronic spray were used to separate and detect the standards at multiple reaction monitoring (MRM) mode. The m^5^dC (m/z 241.9→126.3) and dG (m/z 268.1→152.3) were chosen as parent and child ion pairs for quantitative detection. The CE voltage of both m^5^dC and dG was 15 eV, and the DP voltage was 40 V, respectively. Standard curves of m^5^dC and dG were first graphed and the level of cytosine methylation was calculated as (m^5^dC/dG) x 100%.

### 2.8 RNA-seq to reveal transcriptional response to DNMT3A deficiency

Total RNA was extracted from 10^6^ of DNMT3A KO or WT cells. Oligo(dT) magnetic beads were used to enrich mRNA. CDNA was obtained using Illumina Truseq™ RNA sample prep Kit, and pair-end sequencing (insert size = 300 bp, read length = 150 bp) was performed according to the standard protocol of Novaseq 6000 (Illumina, CA, USA). Raw sequencing reads were filtered to include only high quality reads in downstream analysis: 1) clip adapter sequence from reads, and remove reads with no insertion; 2) clip 3’ low quality bases (Phred quality < 20), and remove the whole read if there exists a single base with Phred quality < 10; 3) remove the reads that have more than 10% ambiguous bases (N); remove the reads that are shorter than 20 bp after clipping. The filtered reads were aligned to human transcriptome (build GRCh38) by TopHat (Trapnell et al., 2009). PCR duplicates were marked and ignored in downstream analysis. All the data were deposited into the open-access Genome Sequence Archive (gsa.big.ac.cn) under accession no. CRA002294.

The read count data of DNMT3A KO and WT cells was analyzed by Cufflink software to identify the differential gene expression induced by DNMT3A deficiency (Trapnell et al., 2012). We used FPKM (Fragments Per Kilobase of exon model per Million mapped reads) to estimate genes expression levels. False discovery rate (FDR) p values were calculated using the method proposed by Benjamini and Hochberg (1995) to correct for multiple testing. Differentially expressed genes in DNMT3A KO cells were identified by FDR p value ≤ 0.05 and absolute logarithm of fold change (log_2_FC) ≥ 2.

### 2.9 KEGG pathway analysis of differentially expressed genes

For the purpose of pathway enrichment analysis, we defined differential expression using a loose definition (FDR p value ≤ 0.05 and absolute log_2_FC ≥ 1). The Ensembl IDs of differentially expressed genes were analyzed by KOBAS (http://kobas.cbi.pku.edu.cn) for KEGG pathway enrichment. The pathways with FDR p value≤ 0.05 were considered significantly differentially expressed.

### 2.10 Bisulfite DNA analysis and quantitative PCR verification of DNMT3A regulated genes

DNMT3A is responsible for the *de novo* methylation of multiple genes, and its mutation can lead to demethylation of promoter CpG and thus elevate gene expression at the transcript level, which further up-regulate or down-regulate related downstream genes indirectly. Therefore, from the gene pool which transcript level was interfered by DNMT3A knockout as determined by RNA-seq, we selected 3 representative genes to verify by bisulfite DNA analysis as well as quantitative PCR: RUNX1, IQGAP3, and DNMT3B. RUNX1 is known to be regulated by DNMT3A in hematopoietic carcinogenesis (Stengel et al., 2018). IQGAP3 is a scaffolding protein that is involved in cancer cells proliferation, and with no correlation with DNA methyltransferases reported before (Lin et al., 2019). All 3 genes were hot studied in malignancy development and helpful to understand the functions of DNMT3A.

DNA methylation status of selected genes were analyzed by bisulfite sequencing PCR (BSP). Genomic DNA was extracted with an Axygen Genomic DNA Miniprep Kit (San Francisco, CA, USA), and 0.5 μg of DNA was modified through bisulfite treatment using a Bisuldream^®^ — Methylation Universal kit (Miozyme; Shanghai, China). Bisulfite-PCR of the genes promoter regions (**Table S1**) was performed using the following specific primers: RUNX1 forward: 5’- TTTTTAGGTTTTAAAATATTTGTGAGTTGT-3’, RUNX1 reverse: 5’- CACCTACCCTCCCCCAAACTATAC-3’, IAGAP3 forward: 5’- GTAGAAAAGGAGTTTGGAAGGAATAAGA-3’, IQGAP3 reverse: 5’- ACTCACAAACTACCCAACCTAAACC-3’, and DNMT3B forward 5’- TTAAAGTAGGATGATAGGTAGGGGTAT-3’, DNMT3B reverse: 5’- CCCTAAAAAATCAAAAACCCTAAAC-3’. The amplified fragments were inserted into pMD19-T vectors (Takara; Tokyo, Japan), and 10-15 clones for each gene were selected for sequencing. The results were analyzed by a web-based quantification tool for methylation analysis (http://quma.cdb.riken.jp).

To detect the transcription levels of the above selected three genes, the DNMT3A KO and WT HEK293 cells were cultured and RNA samples were extracted using Direct-zol RNA kit (Zymo Research; Irvine, CA, USA). Then cDNA was synthesized according to the protocol of the RT-PCR kit (Takara; Kusatsu, Shiga, Japan) and used as templates for quantitative PCR. The primers were designed using Primer Primier 5.0 (Premier Biosoft; Palo Alto, CA, USA) according to published sequences (NCBI Accession number: D43967 for RUNX1, AB105103 for IQGAP3, AF156487 for DNMT3B and M33197 for GAPDH). The following sequences for primers were synthesized (Sangon Biotech; Shanghai, China) as RUNX1 forward: 5’ – TCTCTTCCTCTATCTTCCA– 3’, RUNX1 reverse: 5’–GGTATGTGCTATCTGCTTA–3’; IQGAP3 forward: 5’–GACCACTACCTAACTCAG–3’, IQGAP3 reverse 5’–GCATCATCAACAACTTCTA–3’; DNMT3B forward: 5’- GGCAAGTTCTCCGAGGTCTCTG-3’, DNMT3B reverse: 5’-TGGTACATGGCTTTTCGATAGGA-3’; and GAPDH forward: 5’–CTCTGGTAAAGTGGATATTGT–3’, GAPDH reverse: 5’– GGTGGAATCATATTGGAACA–3’). The real-time PCR procedures were performed with 25 μL PCR reaction systems including 12.5 μL qPCR Mix (Toyobo; Osaka, Japan), 0.4 μM of each primer, and 1 μL template cDNA by thermocycler (StepOnePlus; ThermoFisher, USA). The delta-delta threshold cycle (ΔΔC_T_) method was used to calculate relative copy numbers of targeted genes related to housekeeping gene *GAPDH*.

## 3 RESULTS

### 3.1 Generation of DNMT3A deficient clones of HEK293

Plasmids pX330-sgRNA1 and pX330-sgRNA2 were co-transfected into HEK293 cells. After limiting dilution, the grown clones were selected by PCR using the HEK293-DNMT3A-For and HEK293-DNMT3A-Rev verification primers. We identified one DNMT3A deficient clone from 17 clones, which showed complete disruption of *DNMT3A* gene and designated it as DNMT3A KO (**Figure 2A**). **Figure 2B** shows 137 and 10 bp deletions in the KO A and KO B alleles respectively, leading to complete ablation of *DNMT3A* due to frameshift mutations. Next, we performed western blot to characterize the expression of DNMT3A in the DNMT3A KO clone. As shown in **Figure 2C**, DNMT3A protein expression was completely abrogated in the selected clone, thereby confirming the successful ablation of DNMT3A.

**Figure 2.**
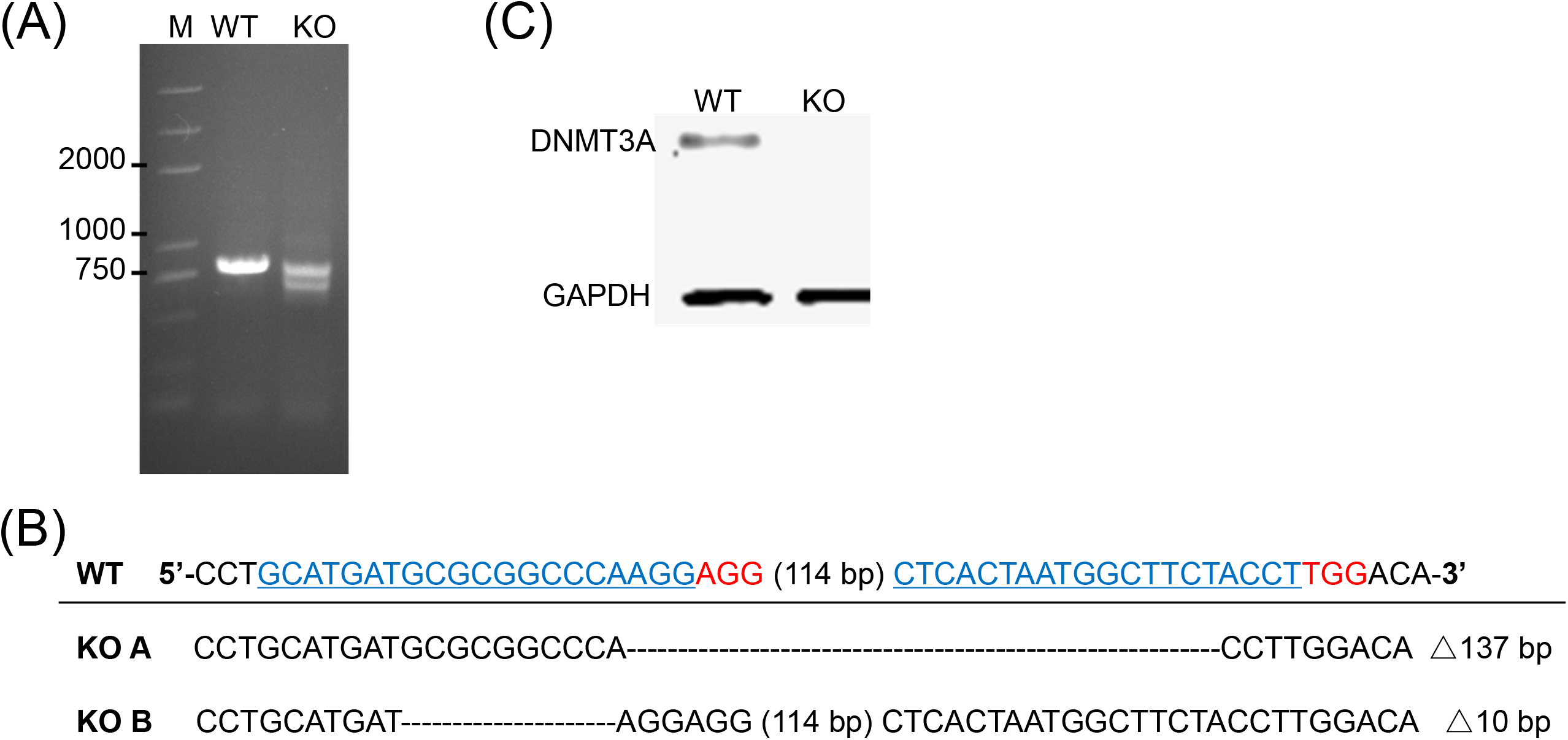
Verification of DNMT3A knockout clone. (A) PCR identification of DNMT3A knockout. Lane M: Trans 2K DNA ladder. (B) Sanger sequencing results of PCR amplicons. KO A and KO B represent the two alleles of *DNMT3A* gene of DNMT3A KO cells. Blue bases: sgRNA sequences; Red bases: PAM. (C) Detection of DNMT3A protein expression with western blot. WT: wild type control, KO: DNMT3A KO cells.

### 3.2 DNMT3A deficiency resulted in genome-wide decrease in DNA methylation

DNMT3A is responsible for the DNA methylation of large number of genes in mammalian cells. To further verify the effect of *DNMT3A*, we performed UPLC-MS to quantify the global DNA methylation level changes following *DNMT3A* knockout. As described in Materials and Methods, we first characterized the peaks of standards A, G, T, C, dA, dG, dC, U, and m^5^dC, and then developed the linear curves of dG and m^5^dC (**Figure S1**). Genomic DNA were extracted from DNMT3A KO and WT cells and hydrolyzed to nucleotides for the measurement of dG and m^5^dC content. The percentage of m^5^dC/dG was calculated to represent the methylation level. As shown in **Figure 3**, the whole-genome DNA methylation level decreased significantly (by 21.5%) in *DNMT3A* KO cells (12.35 ± 0.36%) than in WT cells (9.69 ± 0.13%).

**Figure 3.**
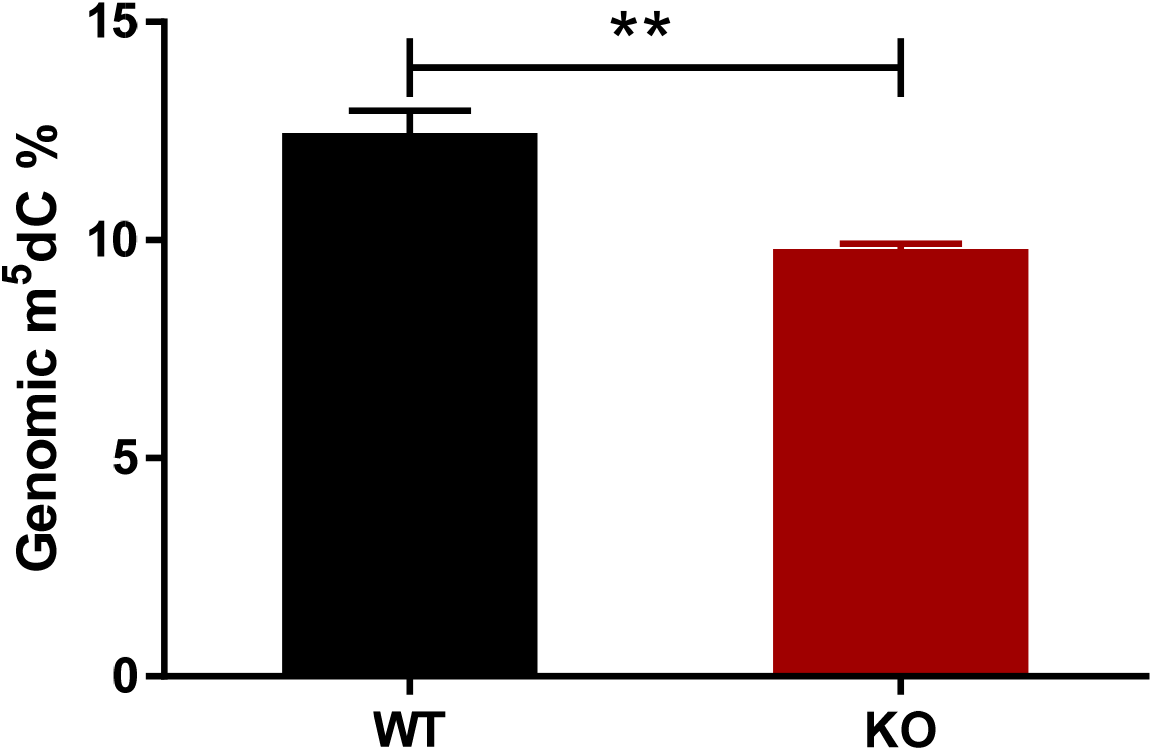
Comparison of methylation level of genomic DNA of wild type (WT) and *DNMT3A* knockout (KO) HEK293 cells. The methylation level of genomic DNA decreased by 21.5% due to DNMT3A deficiency. **p < 0.01 by two tailed students’ t-test.

### 3.3 DNMT3A deficiency impaired cell growth

To evaluate the effect of DNMT3A deficiency on cell proliferation, growth profiles of *DNMT3A* KO and WT cells were evaluated as shown in **Figure 4**. The proliferation ability of HEK293 cells was significantly reduced in response to *DNMT3A* deficiency. After 6 days, the cell counts of *DNMT3A* KO cells reduced to only 40% of WT cells (0.77 ± 0.15) × 10^6^ *vs* (1.94 ± 0.17) × 10^6^ cells). Further, the doubling time was notably prolonged from 0.99 ± 0.28 days for WT cells to 1.53 ± 0.39 days for *DNMT3A* KO cells.

**Figure 4.**
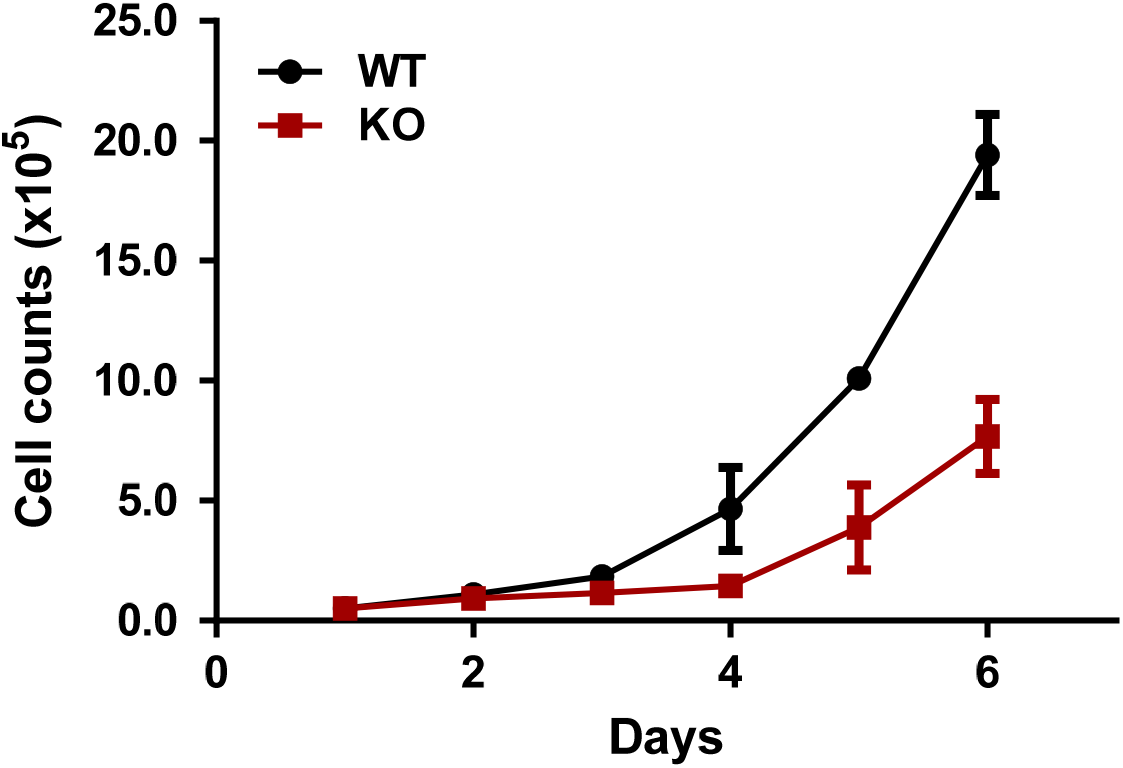
DNMT3A deficiency impaired cell growth. WT: wild type cells; KO: *DNMT3A* KO cells.

### 3.4 RNA-seq analysis

After clipping and filtering, RNA-seq yielded sequencing data of 53.2 million reads (7.9 billion base pairs) and 54.8 million reads (8.2 billion base pairs) for *DNMT3A* KO and WT cells, respectively. It was equivalent to 264.8 and 273.5 times coverage of human transcriptome (30 million base pairs in size). At least 98.4% of the bases had Phred quality > 20 (error rate < 0.01%). TopHat mapped 94.0% of sequencing reads to human genome, including 3.4% reads mapped to multiple genomic position which were excluded from the expression analysis.

### 3.5 Differentially expressed genes and pathways

At significant level of FDR p value ≤ 0.05 and with absolute log_2_FC ≥ 2, we identified 51 differentially expressed genes (**Figure 5**). Among them, more genes were down-regulated (N = 34) as compared to up-regulated genes (N = 17). The top 10 differentially methylated genes are listed in Table 1 (FDR p value ≤ 2.46×10^−42^). The pathway enrichment analysis was performed for 815 up-regulated and 658 down-regulated Ensembl IDs (FDR p value ≤ 0.01 and absolute fold change ≥ 1.5). Pathways related to calcium signaling, ECM-receptor interaction, and Hippo signaling, were up-regulated (**Table 2**), while pathways including Ribosome biogenesis and cysteine and methionine metabolism were down-regulated (FDR p value ≤ 0.01, **Table 3**).

**Table 1.** Highly differentially expressed genes

| Gene ID | Gene Name | FC | log2FC | p-value | p-adjust | regulate |
| --- | --- | --- | --- | --- | --- | --- |
| ENSG00000128591 | FLNC | 0.0363 | -4.78 | 2.17E-175 | 4.00E-171 | down |
| ENSG00000161671 | EMC10 | 0.009 | -6.79 | 4.81E-164 | 4.44E-160 | down |
| ENSG00000019549 | SNAI2 | 0.0632 | -3.98 | 1.35E-97 | 8.32E-94 | down |
| ENSG00000165512 | ZNF22 | 0.0252 | -5.31 | 1.58E-95 | 7.31E-92 | down |
| ENSG00000184368 | MAP7D2 | 0.0452 | -4.47 | 1.99E-80 | 7.35E-77 | down |
| ENSG00000173530 | TNFRSF10D | 0.033 | -4.92 | 2.00E-72 | 6.15E-69 | down |
| ENSG00000164853 | UNCX | 0.0516 | -4.27 | 9.81E-70 | 2.58E-66 | down |
| ENSG00000121413 | ZSCAN18 | 0.0452 | -4.47 | 3.45E-51 | 7.95E-48 | down |
| ENSG00000148798 | INA | 0.0868 | -3.53 | 1.22E-48 | 2.51E-45 | down |
| ENSG00000131435 | PDLIM4 | 0.0543 | -4.20 | 1.34E-45 | 2.46E-42 | down |

**Note:** Gene ID: Ensembl IDs. FC: The fold change of the two samples, and wild type is the control. Log2FC: The base is the base 2 logarithm of the difference between the two samples, and wild type is the control. p-value: The difference in test results of the gene in two samples. p-adjust: Checked result for p-value. Regulate: To indicate whether the expression of the gene is down-regulated or up-regulated; “down” in the column indicates down-regulation of gene expression.

**Table 2.**
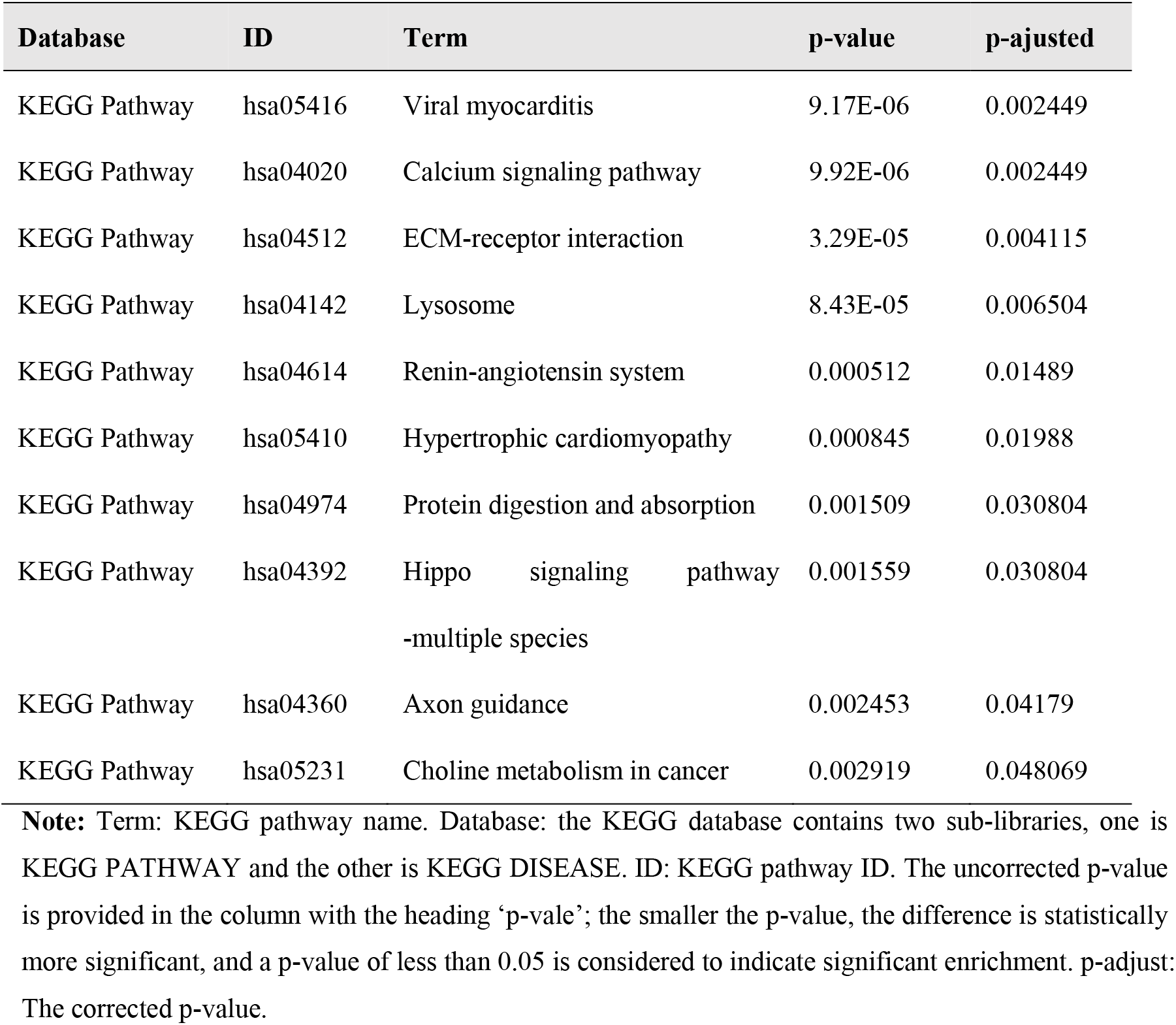
Enrichment analysis of up-regulated pathways

**Table 3.** Enrichment analysis of down-regulated pathways

| Database | ID | Term | p-value | p-adjust |
| --- | --- | --- | --- | --- |
| KEGG Pathway | hsa03008 | Ribosome biogenesis in eukaryotes | 9.9E-07 | 0.0005 |
| KEGG Pathway | hsa00270 | Cysteine and methionine metabolism | 7.4E-05 | 0.0103 |
| KEGG Pathway | hsa05205 | Proteoglycans in cancer | 0.00097 | 0.0854 |
| KEGG Pathway | hsa00670 | One carbon pool by folate | 0.00104 | 0.0854 |
| KEGG Pathway | hsa00230 | Purine metabolism | 0.00258 | 0.1621 |
| KEGG Pathway | hsa05169 | Epstein-Barr virus infection | 0.0027 | 0.1621 |
| KEGG Pathway | hsa03010 | Ribosome | 0.00313 | 0.1621 |

**Note:** Term: KEGG pathway name. Database: the KEGG database contains two sub-libraries, one is KEGG PATHWAY and the other is KEGG DISEASE. ID: KEGG pathway ID. The uncorrected p-value is provided in the column with the heading ‘p-vale’; the smaller the p-value, the difference is statistically more significant, and a p-value of less than 0.05 is indicate significant enrichment. p-adjust: The corrected p-value.

**Figure 5.**
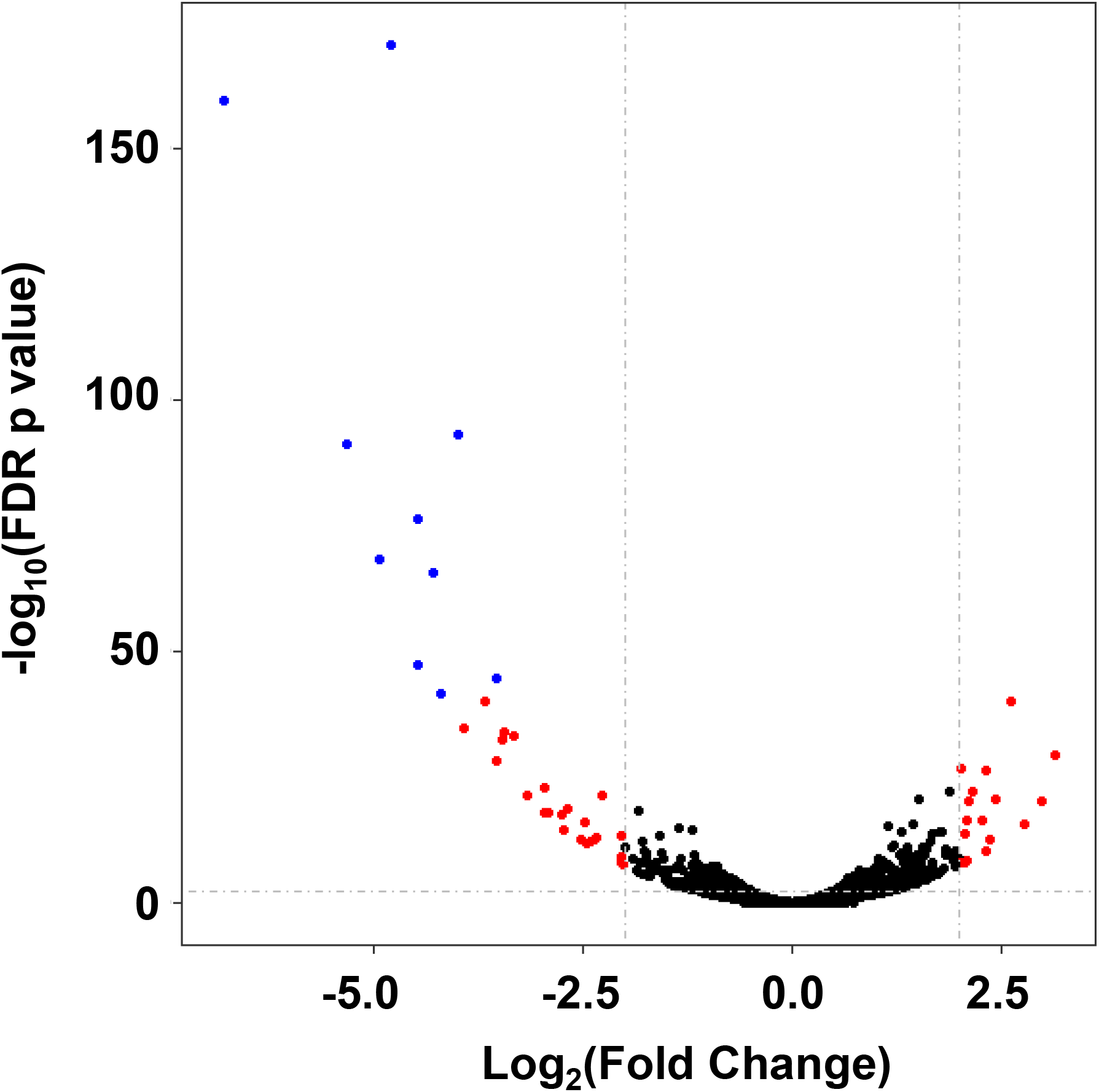
The significance and fold change of differential gene expression induced by *DNMT3A* deficiency. Each dot represents a gene; the significantly differentially expressed genes (FDR p value ≤ 0.05 and fold change ≥ 2) are shown in red; the blue dots are top 10 differentially expressed genes (FDR p value ≤2.46×10^−42^).

### 3.6 Methylation status and transcript level of representative genes regulated by DNMT3A deficiency

According to the results of RNA-seq, there were many up- or down-regulated genes (**Figure 5)**, indicating the alteration in methylation profiles caused by DNMT3A deficiency. Since both DNMT3A and DNMT3B are responsible for *de novo* DNA methylation together, the RNA-seq signal of DNMT3B was determined. We observed a 1.31-fold increase in the RNA-seq signal of DNMT3B in *DNMT3A* KO cells compared to that in WT, indicating the possible compensatory effect of DNMT3B at the deficiency of DNMT3A. Up-regulation of DNMT3B may result in the methylation of some genes and the reduction of their transcription, which explains why the transcription levels of some genes were reduced in this study.

The promoter methylation levels and mRNA transcription levels of *DNMT3B* and two representative tumorigenesis-related genes, *RUNX1* and *IQGAP3*, were verified. The CpG island-rich promoter region (from −1.0 to 0 kb relative to the transcription start site) was analyzed by BSP for each of the three genes. According to the results shown in **Figure 6**, DNMT3A deficiency did reduce the DNA methylation level of *RUNX1* promoter (**Figure 6A**), but it induced the methylation of *IQGAP3* promoter (**Figure 6B**); this induction of methylation was possibly caused by *DNMT3B*, which showed reduced DNA methylation in its promoter region (**Figure 6C**). Quantitative PCR results confirmed the methylation regulation results. Transcription of *RUNX1* was up-regulated by 80%, and that of *IQGAP3* was reduced by 46%. The transcription of *DNMT3B* was also elevated in *DNMT3A* KO cells, but only by 15% (**Figure 6D**). This is the first report to show that DNMT3A contributes to the methylation of the *DNMT3B* gene, indicating the cross-activity of the two d*e novo* DNA methyltransferases.

**Figure 6.**
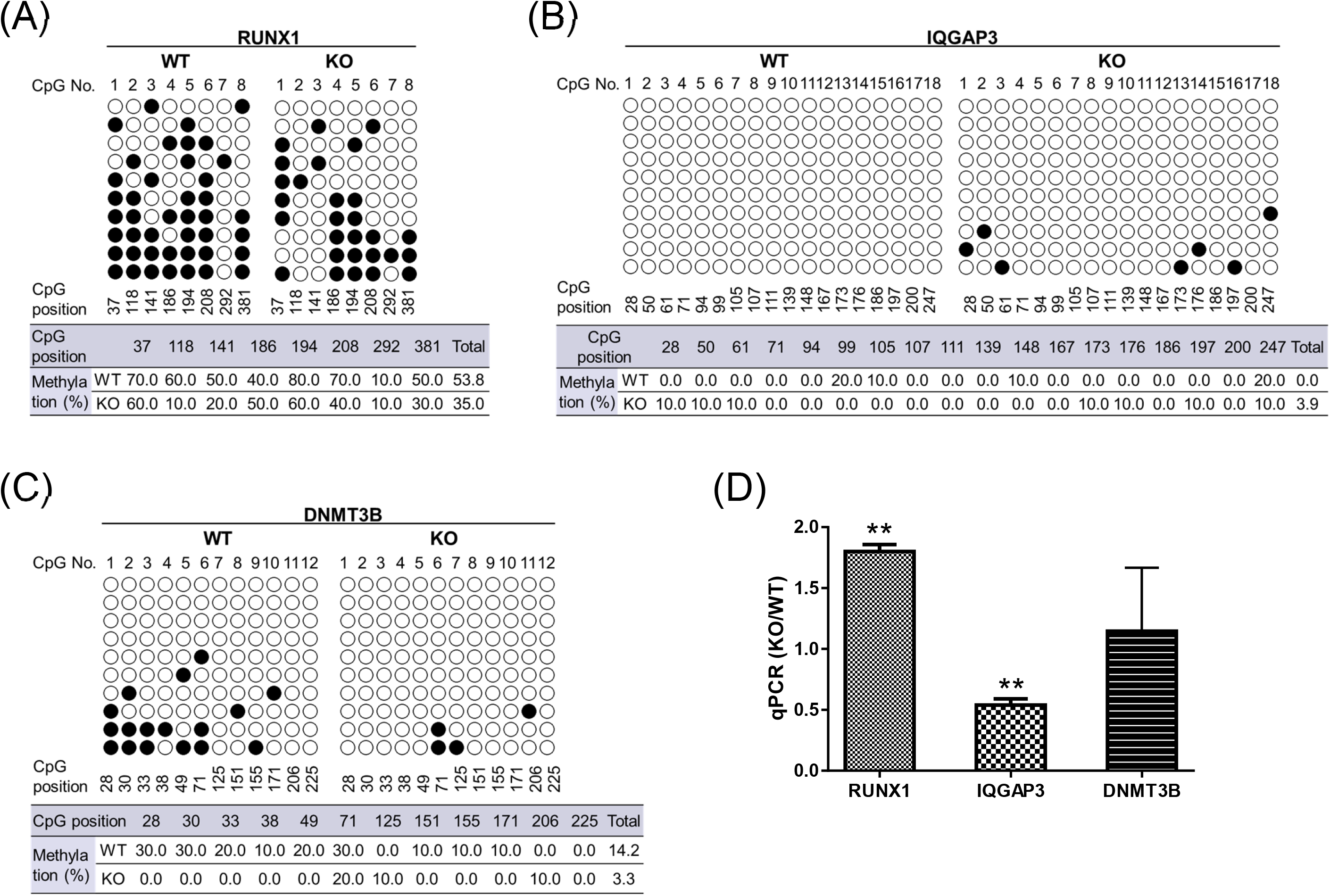
DNA methylation levels and quantitative PCR evaluation of the three representative genes, RUNX1, IQGAP3, and DNMT3B, regulated by DNMT3A deficiency. Bisulfite analysis showed that, in comparison with WT cells, the promoter methylation level of *RUNX1* is decreased by DNMT3A deficiency A), while the *IGAPQ3* promoter has more methylated CpG islands in *DNMT3A* KO cells (B). *DNMT3B* promoter methylation is also regulated by DNMT3A and decreases upon its deficiency (C). (D) Quantitative PCR was performed in triplicate for mRNA expression profiles. In comparison with the relative transcription levels (to *GAPDH* level) in WT cells, *RUNX1* transcription levels were significantly up-regulated by 80%, *IQGAP3* transcription levels were reduced by 46%, and *DNMT3B* transcription levels were up-regulated by about 15%. **p < 0.01 by two tailed students’ t-test.

## 4 DISCUSSION

In recent years, DNMT3A has been intensely studied for its role in tumor prognosis or therapy (Gao et al., 2015; Yang et al., 2016). To better reveal the functions of DNMT3A in cancer occurrence and development, in this study, we mutated HEK293 cells using the CRISPR/Cas9 technology and successfully created a *DNMT3A* knockout cell line, with homozygous frameshift deletion in both alleles. LC-MS analysis showed that knockout of *DNMT3A* induced a 21.5% reduction of global DNA methylation. The reserved DNA methylation could be attributed to the functions of DNMT1 and DNMT3B (Liao et al., 2015). In addition, we attempted the mutation of *DNMT1* or *DNMT3*B using the same strategy in HEK293 cells, but no single clone with the required gene mutation or deficiency was accessed, or the selected clones were unstable for long-term culture (data not shown).

Several previous studies have focused on *DNMT3A* gene knockout in human or mouse-resourced cells, including human embryotic stem cells, human leukemia cells K562, mouse hematopoietic stem cells, as well as mouse somatic cells (Banaszak et al., 2018; Hatazawa et al., 2018; Jeong et al., 2018; Liao et al., 2015). Compared to mouse cells, human cells are less tolerant to DNMT3A deficiency and it can cause lethality and genomic instability in the cells. The results of these previous studies were consistent with our observations that DNMT3A deficiency suppresses HEK293 cell activity. The doubling time of cells dropped from 0.99 ± 0.28 days to 1.53 ± 0.39 days; this result was similar to the phenomenon of impaired cell growth caused by the *DNMT3A* mutation in K562 cells (Banaszak et al., 2018). We assume that this effect is associated with the MAPK or PI3K-Akt pathways, which predominantly contribute to cell proliferation and migration. In this study, we also observed inhibition of the Erk, JNK, and Akt signaling pathways (**Fig. 7**).

**Figure 7.**
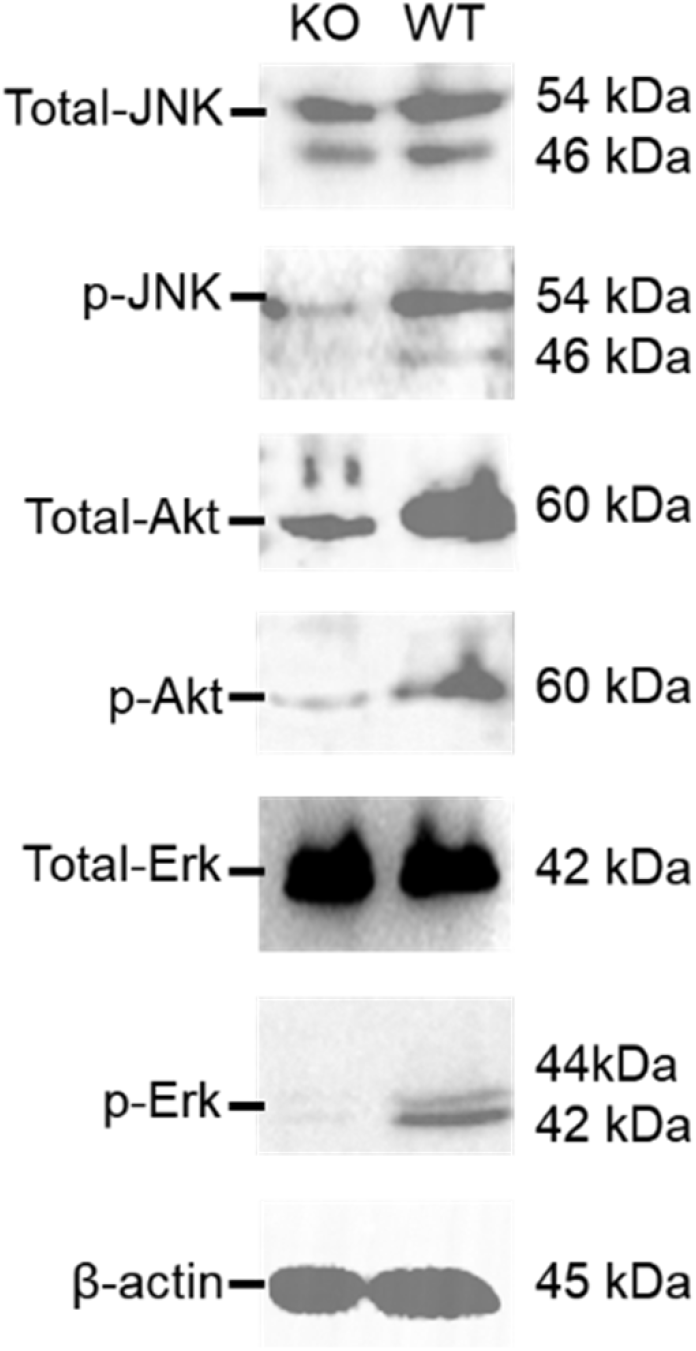
DNMT3A deficiency suppressed MAPK and PI3K-Akt pathways. Total proteins as well as phosphorylated fractions of Erk, JNK, and Akt were detected with β-actin as the housekeeping control.

A previous study introduced frameshift mutations at exons 2 and 3, ablating DNMT3A from more upstream regulatory region (Banaszak et al., 2018). However, in our study, the *DNMT3A* mutation was targeted at exon 19 in the catalytic domain. Reduced genome-wide DNA methylation level in *DNMT3A* KO cells was expected to result in higher transcription levels. However, we unexpectedly observed that a high number of genes were down-regulated in our significant differential expression spectrum with FDR p value ≤ 0.05 and fold change ≥ 2 (binomial p value = 7.6×10^−3^). We also found that the top 10 genes in the most significant gene cluster were down-regulated. DNMT3B showed abnormal up-regulation upon DNMT3A deficiency (**Fig. 6**), and possibly had a methylation function on some of the genes. However, a new research indicates that two SU(VAR)3-9 homologs, the transcriptional anti-silencing factor SUVH1, and SUVH3, as the methyl reader candidates, are associated with euchromatic methylation *in vivo* (Du et al., 2014). In plant, yeast, and mammalian cells, ectopic recruitment of DNAJ1 was shown to enhance gene transcription (Harris et al., 2018). Therefore, the SUVH proteins bind to methylated DNA and recruit DNAJ proteins to enhance proximal gene expression, counteracting the repressive effects of transposon insertion near genes [42]. The top 10 differently expressed genes were likely associated with the SUVH1 and SUVH3 factors when methylation was decreased. However, the real reason for the down-regulation of the top 10 genes in this study is still unknown.

Further investigation of the regulated pathways helped us in understanding that the lower growth rate is a consequence of DNMT3A deficiency. The calcium signaling pathway and ECM-receptor interaction, which are genetically associated with the progression and recurrence of atrial fibrillation (Buttner et al., 2018), and the Hippo signaling pathway were up-regulated in *DNMT3A* KO cells (FDR p values were 0.002449, 0.004114, and 0.03080, respectively). Twist2 is known to regulate ITGA6 and CD44 expression in the ECM-receptor interaction pathway to promote kidney cancer cell proliferation and invasion (Zhang et al., 2016). The major functions of the Hippo pathway are the restriction of tissue growth in adults and modulation of cell proliferation, differentiation, and migration in developing organs (Meng et al., 2016). Cysteine and methionine metabolism are strictly indispensable to the proliferation of porcine adipogenic precursor cells. After commitment, Met deficiency in media has also been shown to affect the differentiation into adipocytes and alter lipid accumulation (Castellano et al., 2017).

In recent years, DNMT3A has been identified to be an ideal target for the development of personalized treatment or predict tumor prognosis (Gao et al., 2015). This is the first report on the effect of DNMT3A disruption in its catalytic domain on genomic DNA methylation and expression. The genes revealed by RNA-seq to be tightly regulated by DNMT3A in HEK293 cells, in this study, are of great significance to understand the functions of DNMT3A in the origin and development of tumors, and are potential novel targets for future cancer therapy.

## ACKNOWLEDGEMENT

This work was funded by the Science and Technology Commission of Shanghai Municipality (No. 17431904500 to Lu H).

## CONFLICT OF INTEREST

The authors declare that they have no conflict of interest.

## ETHICAL APPROVAL

This article does not contain any studies with human participants or animals performed by any of the authors.

## Supplementary Materials

**Table S1.**
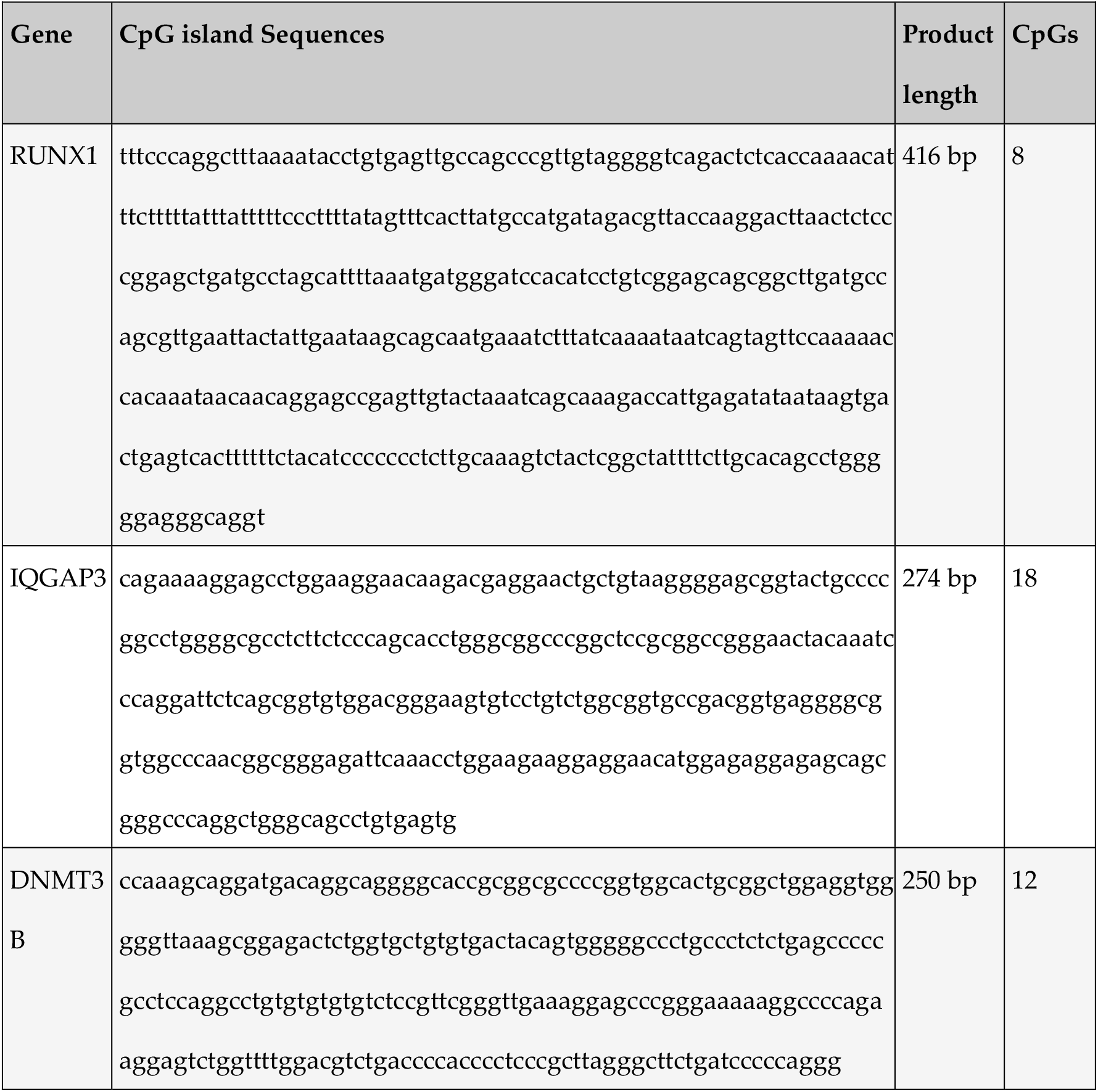
Promoter region sequences and CpG islands of representative genes for bisulfite DNA analysis.

**Figure S1.**
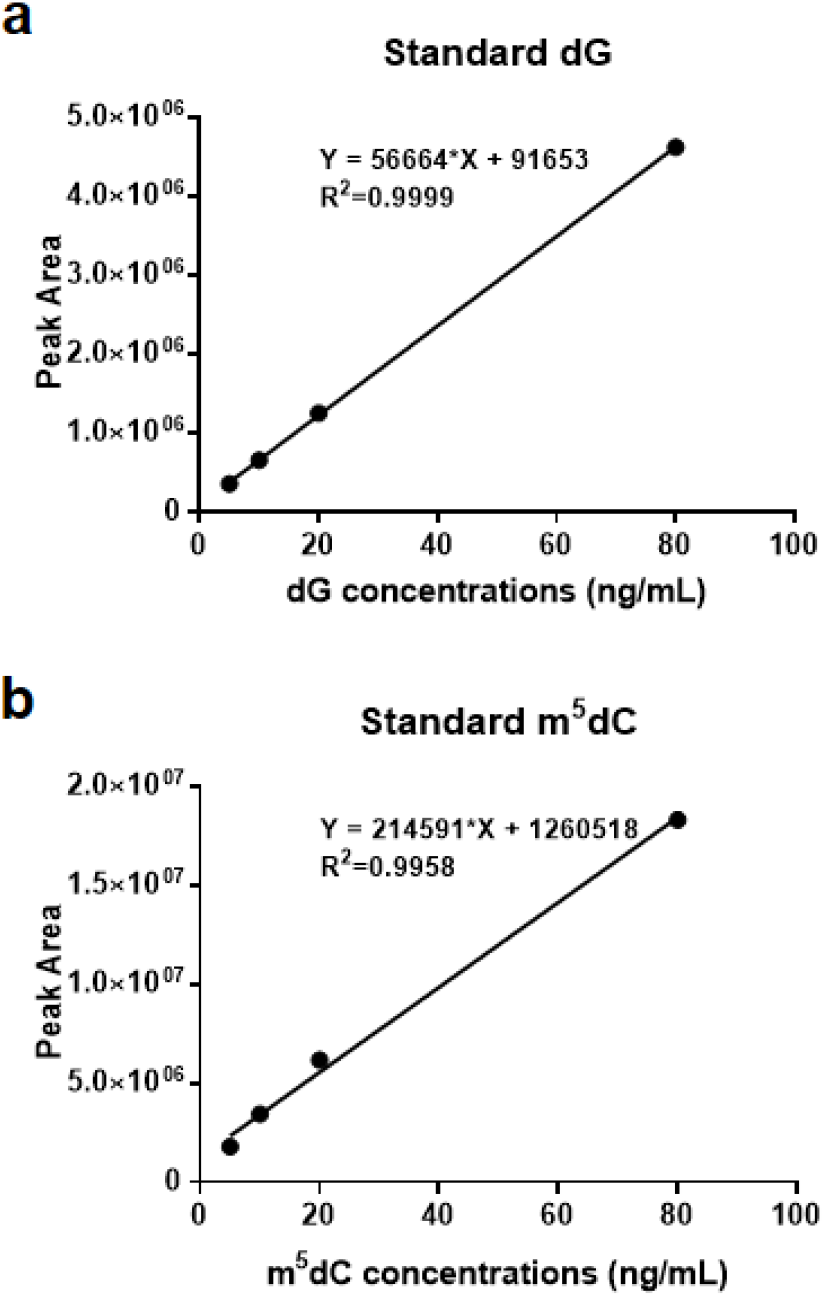
Standard curves of dG and m^5^dC for determination of genomic DNA methylation using UPLC-MS.

## Notes

### Competing Interest Statement

The authors have declared no competing interest.

